# GPI lipid remodeling regulates lipophagy by forming lipid domains in response to glucose deprivation

**DOI:** 10.64898/2026.04.13.718349

**Authors:** Kuya Matsunaga, Kazuki Hanaoka, Yujia Yang, Hinako Nishii, Antonio Cordones-Romero, Sergio Lopez-Martin, Manuel Muñiz, Kouichi Funato

## Abstract

Lipophagy is an important microautophagic process that degrades lipid droplets (LDs) to mobilize stored lipids as an energy source during nutrient starvation. However, the molecular mechanisms regulating lipophagy in response to nutrient starvation remain poorly understood. We found that budding yeast mutants defective in glycosylphosphatidylinositol (GPI) lipid remodeling exhibited aberrant accumulation of lipid droplets (LDs) and neutral lipids under glucose starvation. Our data suggest that the accumulation results from a failure of vacuolar liquid-ordered (Lo) domain-mediated lipophagy. Furthermore, we demonstrated that glycosylphosphatidylinositol-anchored proteins (GPI-APs) localize to vacuoles in response to glucose depletion and that a mutant defective in endocytosis has defects in both vacuolar Lo domain formation and lipophagy. These results imply that GPI lipid remodeling is required for Lo domain-mediated lipophagy upon glucose starvation. We propose that endocytosis functions to supply the lipid portion of GPI-APs, remodeled to C26 diacylglycerol, to the vacuolar membrane for Lo domain formation.

**Summary Statement:** Our data suggest that the endocytic transport of GPI-APs remodeled with C26 diacylglycerol to the vacuole is required for vacuolar Lo domain formation and subsequent lipophagy in response to glucose deprivation. This reveals the essential role of GPI lipid remodeling in ensuring lipophagy to adapt to changes in nutrient availability.

## Introduction

Lipid droplets (LDs) are cellular organelles that store and supply neutral lipids, primarily triacylglycerols (TAGs) and sterol esters (SEs), and play a central role in cellular lipid and energy homeostasis. Cells dynamically regulate the biosynthesis and degradation of LDs to adapt to changes in nutrient availability and environmental conditions, and impairments in these processes are associated with metabolic disorders such as non-alcoholic fatty liver disease (NAFLD) and metabolic syndrome (Wang, 2016; Rahman et al., 2021; Schott et al., 2022; Zhang et al., 2022). Lipids stored in LDs are primarily degraded through two pathways: lipase-mediated lipolysis and lipophagy, a selective form of microautophagy. When nutrients such as glucose, the main carbon source, are depleted, cells induce lipophagy to mobilize stored lipids as energy sources (van Zutphen et al., 2014; Wang, 2016; Rahman et al., 2021; Schott et al., 2022). However, the molecular mechanisms that govern and promote lipophagy in response to nutrient starvation remain unclear.

We have previously reported that the biosynthesis of glycosylphosphatidylinositol (GPI), a glycolipid used in the post-translational modification of many cell surface proteins, and GPI anchoring to proteins are involved in regulating LD biogenesis in the budding yeast *Saccharomyces cerevisiae* (Kajiwara et al., 2008). GPIs consist of acyl-phosphatidylinositol (acyl-PI)-based lipid moieties and glycan moieties containing glucosamine, mannose, and ethanolamine phosphate (Muñiz and Riezman, 2016; Kinoshita and Fujita, 2016). After GPI synthesis in the endoplasmic reticulum (ER) lumen, the attachment of GPI to proteins is catalyzed by the GPI transamidase complex. The resulting GPI-anchored proteins (GPI-APs) undergo stepwise remodeling of their lipid moiety (Muñiz and Riezman, 2016; Kinoshita and Fujita, 2016). In *S. cerevisiae*, Bst1 removes the acyl chain from inositol (Tanaka et al., 2004), followed by the removal of the sn-2 unsaturated fatty acid on diacylglycerol by Per1 (Fujita et al., 2006), resulting in the generation of lyso-GPI. Subsequently, Gup1 adds a C26 very-long-chain saturated fatty acid to form a C26 diacylglycerol moiety (referred to as GPI lipid remodeling) (Bosson et al., 2006). Furthermore, in some cases, the C26 diacylglycerol generated is exchanged with C26 very long-chain saturated ceramide by Cwh43, yielding C26 inositolphosphoceramide (IPC), a process termed ceramide remodeling (Umemura et al., 2007; Ghugtyal et al., 2007). As Per1 and Gup1 are essential for the segregation of GPI-APs into detergent-resistant membranes (DRMs) (Fujita et al., 2006; Bosson et al., 2006; Umemura et al., 2007), it has been proposed that the C26 diacylglycerol moiety of GPI assembles GPI-APs into liquid-ordered (Lo) lipid domains on the ER membrane, and that GPI-APs are transported to the cell surface via a lipid domain-dependent secretory pathway (Muñiz and Riezman, 2016).

GPIs are increasingly recognized not only as membrane anchors and determinants of protein sorting but also as regulators of various intracellular processes, including cell cycle progression, Golgi and vacuole biogenesis, and endocytosis (Fujita et al., 2004; Chen, L et al., 2021; Rodriguez-Gallardo et al., 2022; Sasaki et al., 2024; Chen and Banfield, 2024; Ye et al., 2025). A crosstalk between GPI biosynthesis and filamentation signaling also exists (Komath, 2024). In *S. cerevisiae*, defects in GPI biosynthesis or anchoring increase the number of LD per cell (Kajiwara et al., 2008). Notably, although it remains unclear whether the *CWH43* homologue of the fission yeast *Schizosaccharomyces pombe* catalyses ceramide remodeling, mutations in *S. pombe CWH43* have been reported to cause significant increases in LD number and neutral lipid levels (Nakazawa et al., 2018). These findings suggest that GPI anchoring and subsequent GPI lipid remodeling are involved in regulating LD biogenesis; however, the underlying mechanisms remain unknown.

In this study, we found that mutants with defects in GPI lipid remodeling caused aberrant accumulation of LDs and neutral lipids under glucose starvation in *S. cerevisiae*. We also revealed that remodeling mutants failed to form vacuolar Lo domains during glucose starvation, leading to impaired lipophagy. Furthermore, we demonstrated that GPI-APs localize to the vacuole in response to glucose depletion and that the *end3*Δ mutant, which is defective in endocytosis, exhibits defects in both Lo domain formation and lipophagy, implying that endocytic trafficking towards the vacuole is a supply route for GPI-APs required for lipophagy in response to nutrient starvation. Thus, our study reveals a regulatory mechanism of lipophagy mediated by the endocytic transport of remodeled GPI-APs to the vacuole.

## Results

### Defects in GPI lipid remodeling increase LD numbers and neutral lipid levels during glucose starvation

Previous studies have shown a link between GPI-AP structure and LD homeostasis (Kajiwara et al., 2008; Nakazawa et al., 2018). To investigate whether GPI lipid remodeling affects LD homeostasis, we analyzed the number of LDs in the *bst1*Δ, *per1*Δ, and *gup1*Δ strains using fluorescence microscopy. Cells were grown in glucose-containing SD medium and either.stained with AutoDot directly or after a 24 h shift to SD medium without glucose (SD-Glucose) to visualize the LDs. The number of LDs per cell was comparable between wild-type (WT) and mutant cells in the glucose-containing medium (Fig. 1A and B). In contrast, in glucose-depleted medium, the mutant strains exhibited a marked increase in the number of LDs compared to WT. Next, we quantified the levels of intracellular neutral lipids, TAG and SE, which are mainly stored in LDs. Neutral lipid levels were increased 1.5-to 2-fold in the mutant strains compared to that in the WT in glucose-containing media (Fig. 1C and D). Under glucose starvation, both SE and TAG levels were elevated 2-to 3-fold in the mutant strains. Ergosterol levels were not affected by the presence or absence of glucose or GPI remodeling mutations. Collectively, these results suggest that GPI lipid remodeling is critical for the regulation of LD number and neutral lipid metabolism during glucose starvation.

**Figure 1.**
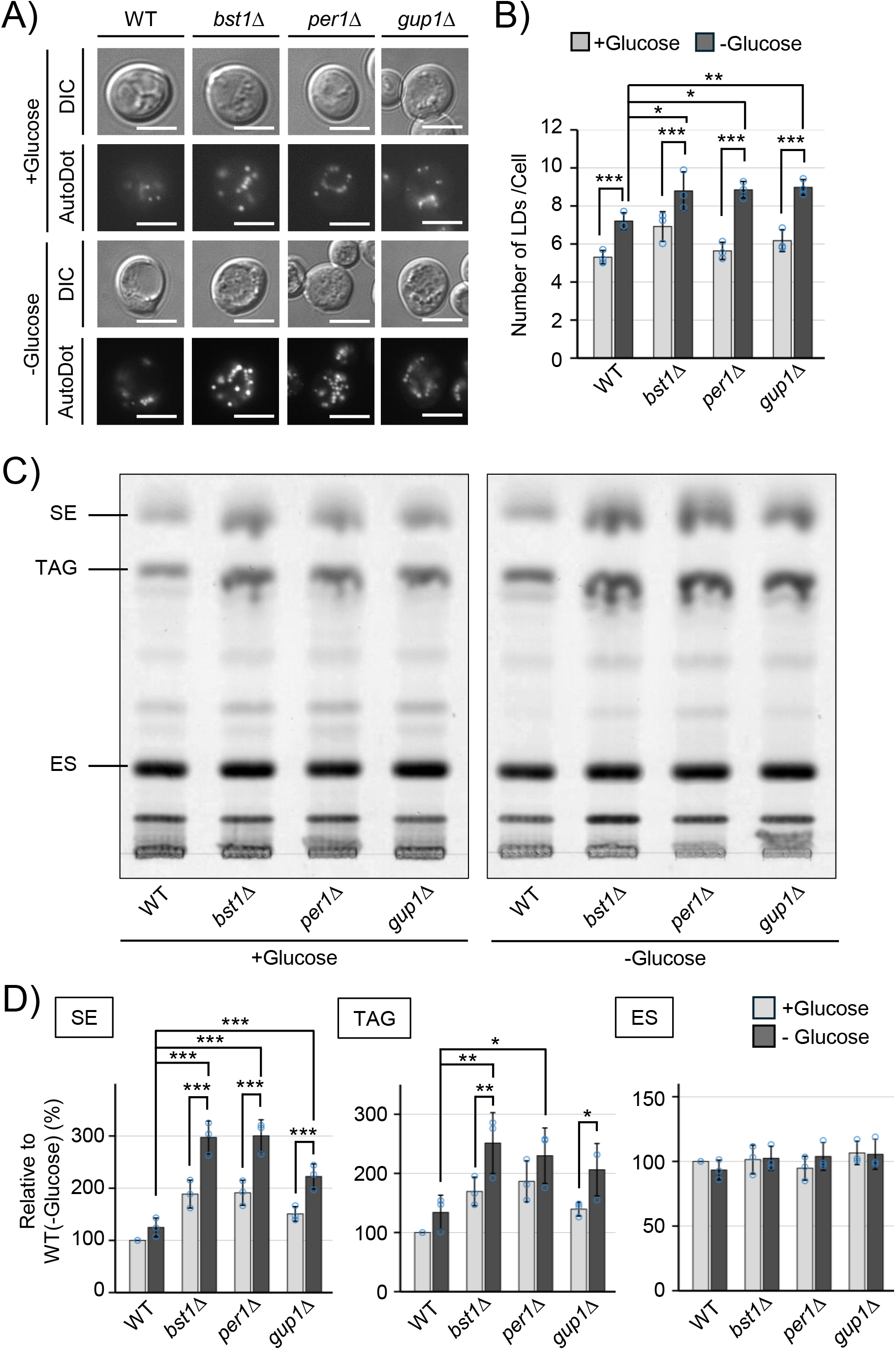
Defects in GPI lipid remodeling result in increased lipid droplet number and neutral lipid levels under glucose starvation. (A), Wild-type and GPI lipid remodeling mutant cells (*bst1*Δ, *per1*Δ, and *gup1*Δ) were grown in SD medium and either analyzed directly or after a 24 h shift to SD-Glucose. Cells were stained with AutoDot and visualized by fluorescence microscopy. Representative images are shown. Scale bar, 5 µm. (B), LD number per cell was quantified. ≥100 cells were analyzed per biological replicate. Data represent mean ± s.d. from three independent experiments (n=3). Statistical significance was determined by one-way ANOVA followed by Tukey’s multiple comparison test. *p<0.05, **p<0.01, ***p<0.001. (C), Neutral lipids were extracted from cells cultured as described in (A) and separated by thin-layer chromatography (TLC). Representative images from three independent experiments are shown. SE, sterol esters; TAG, triacylglycerol; Erg, ergosterol. (D) The relative abundance of each lipid species was quantified. Levels in WT cells grown in SD medium were set to 100. Data represent mean ± s.d. from three independent experiments (n=3). Statistical significance was determined by one-way ANOVA followed by Tukey’s multiple comparison test. *p<0.05, **p<0.01, ***p<0.001.

### GPI lipid remodeling is required for LD degradation via lipophagy during glucose starvation

LDs accumulated during nutrient starvation are degraded via lipophagy, which is a form of microautophagy (Fei et al., 2008; van Zutphen et al., 2014; Wamg et al., 2014). To determine whether the LD accumulation observed in GPI lipid remodeling mutants results from impaired lipophagy, we assessed the cleavage of Erg6-GFP, a lipophagy reporter. Similar to previous studies (Wang et al., 2014; Seo et al., 2017), free GFP was not detected in glucose-containing medium. Upon glucose starvation, while free GFP was generated in WT cells, it was significantly reduced in the GPI lipid remodeling mutants (Fig. 2A and B), indicating that Erg6-GFP degradation was impaired in the mutant cells. The degradation defects were not due to impaired transport or activity of vacuolar proteases, given that vacuolar carboxypeptidase Y (CPY) has been reported to undergoes normal maturation in *bst1*Δ, *per1*Δ, and *gup1*Δ cells (Bosson et al., 2006; Fujita et al., 2006; Rodriguez-Gallardo et al., 2022). Furthermore, unlike the *vma3*Δ mutant, which has a defect in vacuolar acidification, these GPI remodeling mutants grew normally at pH 7.5 (Fig. S1), implying that vacuolar acidification, which is a prerequisite for protease activation, occurred normally. Collectively, these results suggest that the reduced Erg6-GFP degradation in the GPI lipid remodeling mutants reflects a lipophagy defect rather than a general vacuolar dysfunction.

**Figure 2.**
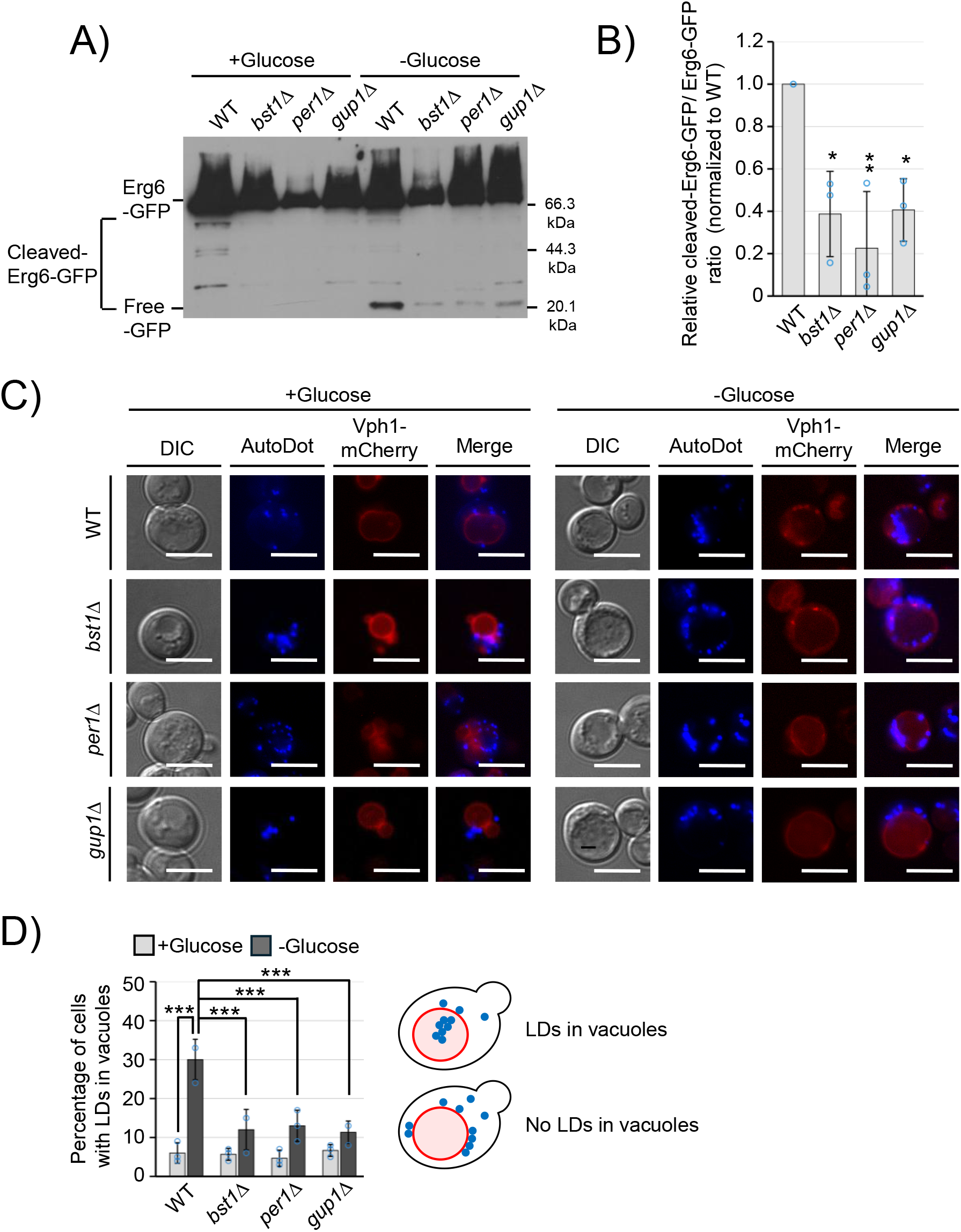
GPI lipid remodeling is required for lipophagy under glucose starvation. , Cells expressing Erg6-GFP, a lipophagy reporter were cultured as in Fig. 1A. Cell extracts were analyzed by Western blotting using anti-GFP antibodies. (B), Erg6-GFP degradation was quantified as the ratio of cleavaged Erg6-GFP to total Erg6-GFP signal. Data represent mean ± s.d. from three independent experiments (n=3). Statistical significance was determined by one-way ANOVA followed by Tukey’s multiple comparison test. *p<0.05, **p<0.01, ***p<0.001. (C), Wild-type and GPI lipid remodeling mutant cells expressing the vacuolar marker Vph1-mCherry were stained with AutoDot, and visualized by fluorescence microscopy. Representative images are shown. Scale bar, 5 µm. (D), The percentage of cells containing LD dot within vacuoles was quantified. ≥100 cells were analyzed per biological replicate. Data represent mean ± s.d. from three independent experiments (n=3). Statistical significance was determined by one-way ANOVA followed by Tukey’s multiple comparison test. ***p<0.001.

To identify which step of lipophagy was impaired in the mutants, we visualized LDs and the vacuoles using AutoDot and the vacuolar marker Vph1-mCherry, respectively. Consistent with the results of the Erg6-GFP cleavage assay, few LDs were observed within the vacuoles before glucose starvation (Fig. 2C and D). Upon glucose starvation, the proportion of cells exhibiting LD internalization into the vacuole was markedly reduced in the GPI lipid remodeling mutants, and many LDs clustered on the cytoplasmic leaflet of the vacuolar membrane. Taken together, these findings suggest that the internalization of LDs into vacuoles during lipophagy is impaired in GPI lipid remodeling mutants.

### GPI lipid remodeling is required for vacuolar Lo domain formation

It has been reported that under glucose starvation, vacuolar membrane segregates into two distinct domains: the liquid-ordered (Lo) domain, enriched in sterols and sphingolipids, and the liquid-disordered (Ld) domain (Toulmay and Prinz, 2013; Numrich et al., 2015; Sakuragi et al., 2023; Kim and Budin, 2024; Juarez-Contreras et al., 2025). During lipophagy, Lo domains act as specific sites on the vacuolar surface that anchor and engulf LDs (Wang et al., 2014). As GPI lipid remodeling has been shown to play a role not only in the segregation of GPI-AP into lipid domains but also in the stabilization and maintenance of lipid domains (Ferreira and Lucas, 2008; Tulha et al., 2008), we hypothesized that remodeled GPI lipid moieties are required for the formation of vacuolar Lo domains. To test this hypothesis, we investigated whether GPI lipid remodeling mutants affect the formation of vacuolar Lo domains upon glucose starvation. We used Ego1-GFP as a marker for the Lo domain (Fig. 3A), and classified vacuolar domain patterns into three types: Type A (no domains), showing a homogeneous distribution; Type B (partial domains), displaying fragmented signals with small gaps; and Type C (coalesced domains), containing large patches lacking Ego1-GFP (Fig. 3B). In the presence of glucose, Type A accounted for more than 90% of the population in both WT and GPI lipid remodeling mutant strains (Fig. 3A and C). In contrast, under glucose starvation, approximately 60% of WT cells showed Type B or C patterns, whereas only approximately 30% of GPI remodeling-deficient cells exhibited these patterns. These results suggest that GPI lipid remodeling is required for efficient vacuolar Lo domain formation.

**Figure 3.**
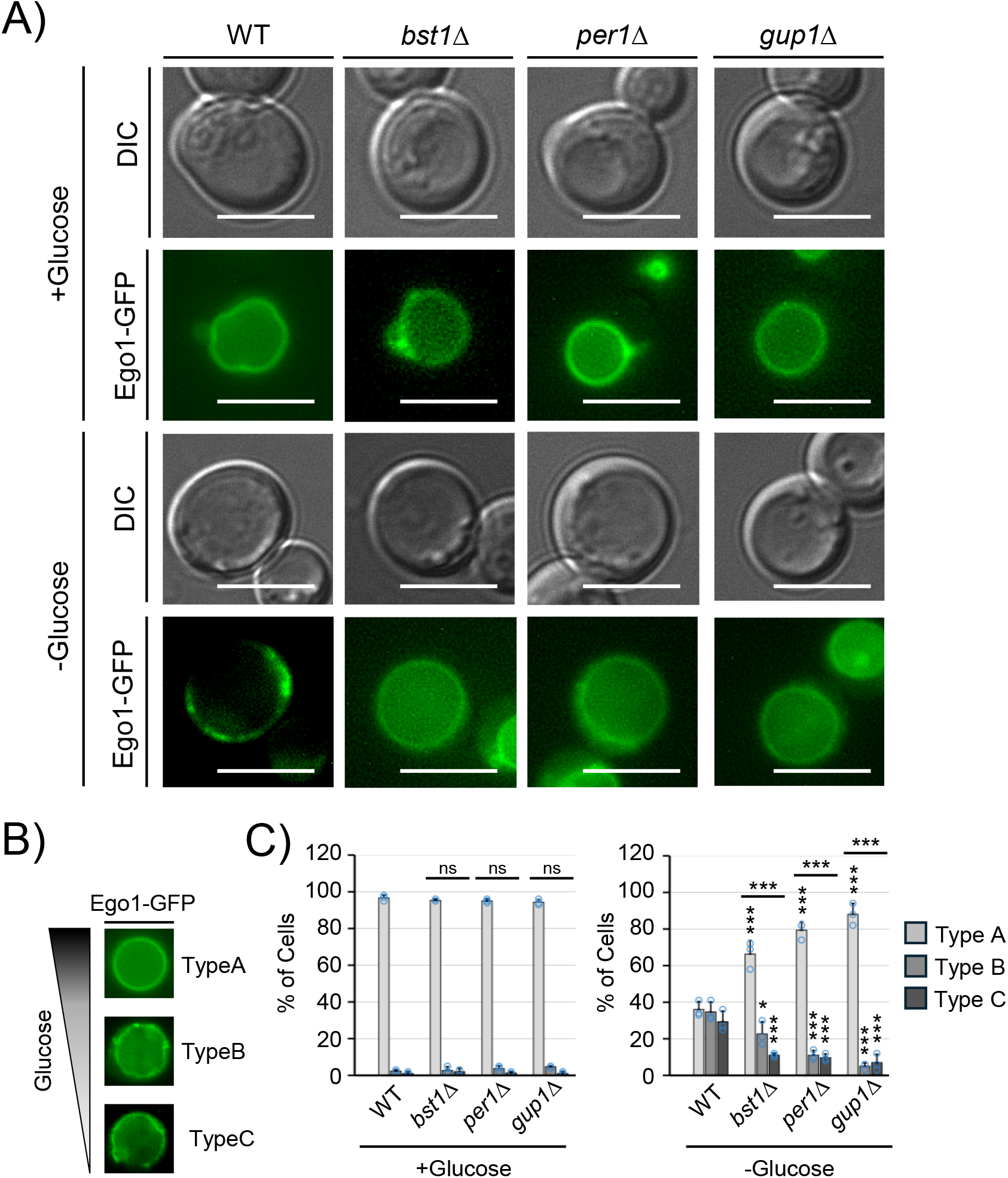
Mutants defective in GPI lipid remodeling impair vacuolar Lo domain formation. (A-C) Wild-type and GPI lipid remodeling mutant cells expressing Ego1-GFP were visualized by fluorescence microscopy. Scale bar, 5 µm (A). Vacuoles were categorized into three types based on Ego1-GFP localization (B): Type A, no Lo domains; Type B, partial domains; and Type C, coalesced domains. Representative examples are shown. Quantification of Lo domain categories. ≥100 cells were analyzed per biological replicate (C). Data represent mean ± s.d. from three independent experiments (n=3). Overall distribution differences between WT and mutants were assessed using a Chi-square test. Comparisons of the same category among strains were performed by one-way ANOVA followed by Tukey’s multiple comparison test. *p<0.05, ***p<0.001.

### Endocytic transport of GPI-APs is required for vacuolar Lo domain formation and lipophagy

Our results suggest that GPI remodeling is involved in the formation of vacuolar Lo domains; however, the underlying mechanism remains unclear. Glucose starvation, which induces vacuolar membrane phase separation, is known to trigger endocytic internalization of various proteins (Laidlaw et al., 2021). Although GPI-APs primarily function at the cell surface, some are often observed to be localized within vacuoles (Fujita et al., 2006; Castillon et al., 2011; MacDonald et al., 2015). We also observed that some Gas1-GFP, a representative GPI-AP present in the plasma membrane, localized to the vacuoles upon a shift from nutrient-rich (YPD) to minimal (SC) media (Fig. S3). Thus, we hypothesized that glucose starvation triggers the endocytic internalization of GPI-APs and delivers them to the vacuole, leading to the formation of vacuolar Lo domains. To test this, we monitored the intracellular distribution of Gas1-GFP after shifting the cells from YPD medium to glucose starvation medium. Gas1-GFP localization was categorized into three patterns based on the relative fluorescence intensity between the plasma membrane (PM) and the vacuole (PM > vacuole, PM = vacuole, or PM < vacuole). In WT cells, the proportion of cells exhibiting stronger GFP signals in the vacuole relative to the plasma membrane (PM < vacuole) increased in a time-dependent manner after the shift to starvation conditions (Fig. 4A and B). In contrast, in the *end3*Δ strain, a mutant defective in endocytosis (Benedetti et al., 1994; Tang et al., 1997), a high proportion of cells exhibited weak vacuolar GFP signals (PM > vacuole). These results indicate that GPI-APs are delivered to the vacuole via endocytosis in response to glucose starvation.

If the delivery of remodeled GPI-APs to the vacuole via endocytosis is essential for vacuolar Lo domain formation, *end3*Δ cells would be defective in Lo domain formation and lipophagy. Indeed, in the *end3*Δ strain, both vacuolar Lo domain formation and Erg6-GFP degradation were significantly impaired (Fig. 4C-F). Collectively, these results suggest that the delivery of remodeled GPI-APs to the vacuole is required for vacuolar Lo domain formation and lipophagy.

**Figure 4.**
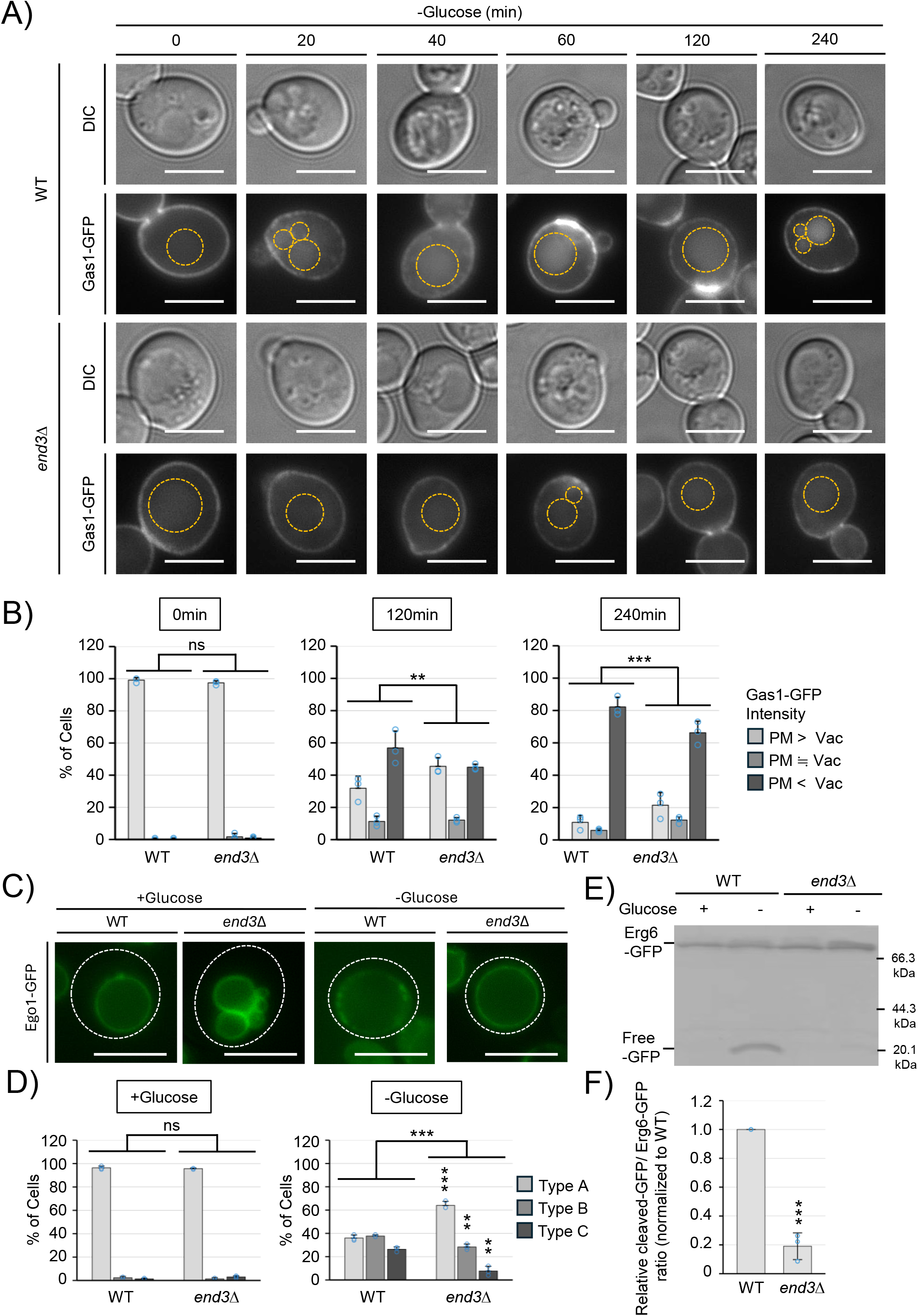
Gas1-GFP is transported to the vacuole via endocytosis during glucose starvation. (A), WT and *end3*Δ cells expressing Gas1-GFP were grown in YPD and either analyzed directly or shifted to SD-Glucose and analyzed at 20, 40, 60, 120, and 240 min by fluorescence microscopy. Representative images are shown. Scale bar, 5 µm., Gas1-GFP localization was classified into three categories based on relative fluorescence intensity between the plasma membrane (PM) and vacuole: PM > vacuole, PM = vacuole, and PM < vacuole. The percentage of cells in each category was quantified at YPD, 120, and 240 min after shifting to SD-Glucose. ≥100 cells were analyzed per biological replicate. Data represent mean ± S.D. from three independent experiments (n=3). Statistical significance was assessed using a Chi-square test. *p<0.05, **p<0.01, ***p<0.001. (C), WT and *end3*Δ were grown in SD medium and either analyzed directly or after a 24 h shift to SD-Glucose and visualized by fluorescence microscopy. (D) Lo domain morphology was classified as described in Fig. 3B. ≥100 cells were analyzed per biological replicate. Data represent mean ± s.d. from three independent experiments (n=3). Differences in overall distribution between WT and *end3*Δ were assessed using a Chi-square test, and individual categories were compared using Student’s t test. **p<0.01, ***p<0.001. (E) WT and *end3*Δ were grown in SD medium and either analyzed directly or after a 72h shift to SD-Glucose. Cell extracts were analyzed by Western blotting using anti-GFP antibodies. (F) Erg6-GFP degradation was quantified as the ratio of free GFP to total Erg6-GFP signal. Data represent mean ± s.d. from three independent experiments (n=3). Statistical significance was determined by one-way ANOVA followed by Tukey’s multiple comparison test. ***p<0.001.

**Figure 5.**
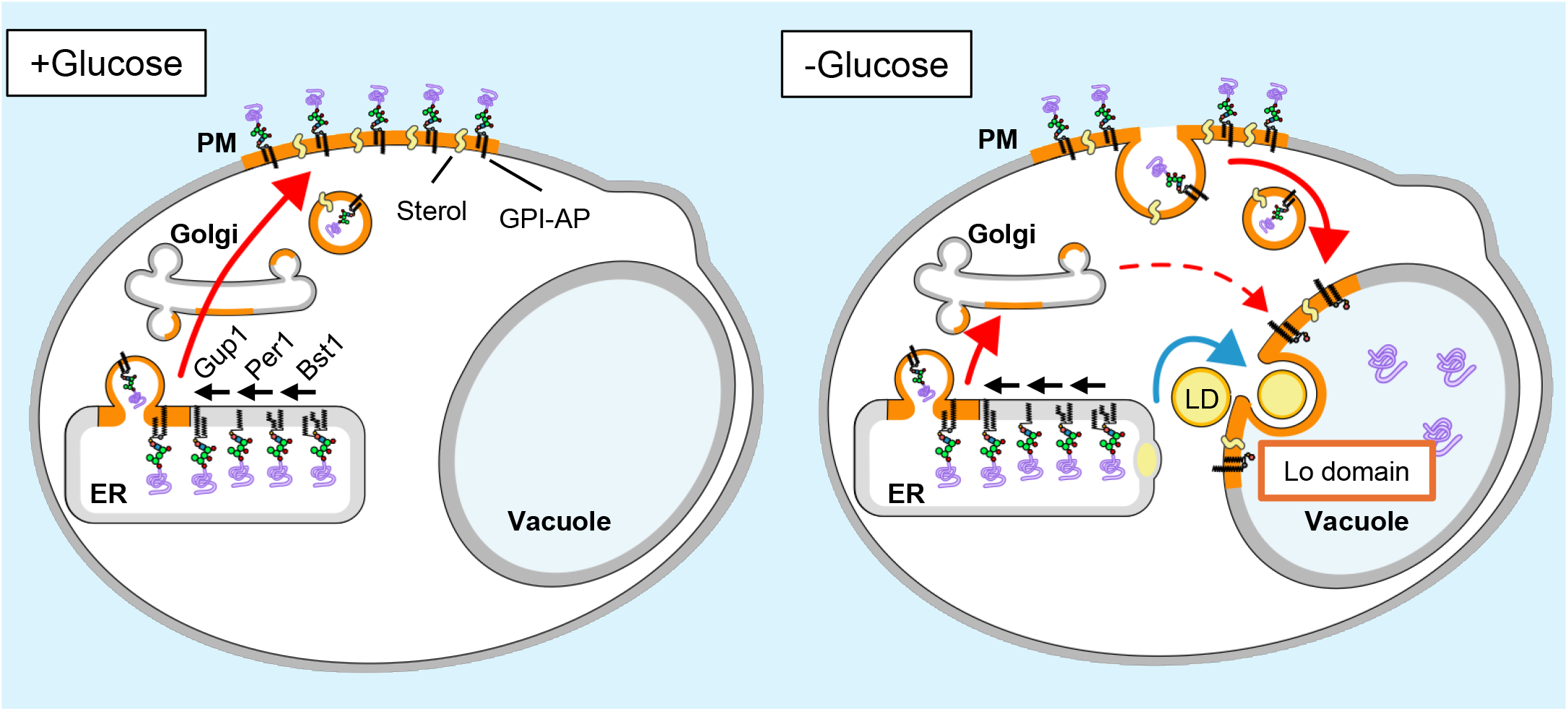
Model for GPI lipid remodeling-dependent regulation of lipophagy through vacuolar membrane domain formation under glucose starvation. Left panel (Glucose-rich conditions). GPI-APs are synthesized and their lipid moiety are remodeled in the ER by Bst1, Per1, and Gup1. The remodeled GPI-APs are transported from the ER to the plasma membrane via vesicular transport (red arrow). Right panel (Glucose starvation). Upon glucose starvation, GPI-APs, which are localized to the plasma membrane, are internalized via endocytosis and delivered to the vacuole (red arrow). The lipid moiety of GPI-APs remains associated with vacuolar liquid-ordered (Lo) domains, whereas the protein moiety is cleaved and released into the vacuolar lumen. LDs accumulate on the cytoplasmic leaflet of vacuolar Lo domains, which serve as platforms for subsequent LD internalization (blue arrow). ER, endoplasmic reticulum; LD, lipid droplet; Lo domain, liquid-ordered membrane domain.

## Discussion

In this study, we demonstrated that GPI lipid remodeling is crucial for vacuolar Lo domain formation and lipophagy during glucose starvation in yeast. We also revealed that endocytic trafficking towards the vacuole serves as a supply route for GPI-APs. Previous studies have proposed that endocytic transport to the vacuole in response to glucose starvation functions in degradation of unnecessary plasma membrane proteins to recycle amino acids and in reorganization of the plasma membrane (Laidlaw et al., 2021). Here, we propose that endocytosis during glucose starvation also functions to supply neutral lipids from LDs via GPI lipid remodeling-dependent lipophagy.

GPI lipid remodeling mutants exhibited defects in Lo domain formation. It was possible that the defects could result from either inefficient transport of unremodeled GPI-APs to the vacuole or the inability of unremodeled GPI-APs to form vacuolar Lo domains. A previous study reported that the internalization of Snc1, an endocytic marker, is impaired in *bst1*Δ and *per1*Δ mutants (Burston et al., 2009). In contrast, another report indicated that tryptophan permease Tat2, a lipid domain-associated transmembrane protein that localizes to the plasma membrane, is mislocalized to the vacuole in *per1*Δ and *gup1*Δ mutants (Yoko-O et al., 2013). We observed that Gas1-GFP accumulated in the vacuoles of *bst1*Δ cells immediately after glucose depletion, and the level of accumulation was higher than that in wild-type cells (Figure S4). This suggests that defects in GPI lipid remodeling do not inhibit endocytosis but rather enhance the trafficking of GPI-APs to the vacuole. We speculate that unremodeled GPI-APs that fail to assemble into lipid domains of the plasma membrane may be endocytosed more rapidly in response to glucose starvation. Consistent with this model, it has been shown that although distinct from the domains where GPI-APs assemble, MCC/eisosomes, which have the characteristics of lipid domains, function to inhibit the endocytosis of transport proteins such as Can1 and Fur4, and disruption of the domain assembly by deletion of Pil1, a component of MCC/eisosomes, leads to an accelerated endocytosis of transporters (Grossmann et al., 2008; Gournas et al., 2018). Similarly, in *S. pombe*, under nutrient-limiting conditions, a plasma membrane-localizing amino acid transporter, Gap1 permease, which fractionates with DRMs, is endocytosed and transported to the vacuole (Nakase et al., 2010). Importantly, despite GPI-AP transport to the vacuole, no Lo domains were formed in *bst1*Δ cells. Taken together, we conclude that the failure of Lo domain formation in GPI lipid remodeling mutants arises from the unremodeled lipid moiety of GPI-APs.

Gas1 remains anchored to the plasma membrane, whereas Cwp2 is transferred and linked to the cell wall (van der Vaart et al., 1995). We observed that, like Gas1-GFP, Cwp2-VENUS also localized to the vacuole under nutrient-poor conditions (Figure S3). Cwp2-VENUS may be transported from the Golgi apparatus or plasma membrane to the vacuole before being incorporated into the cell wall. This raises the possibility that in response to environmental stress, cells mobilize as many available GPI-APs as possible to assemble vacuolar lipid domains for energy production via lipophagy.

The biosynthesis of sphingolipids and sterols and their delivery to the vacuole, likely occurring at the membrane contact site, are essential for vacuolar Lo domain formation (Toulmay and Prinz, 2013; Numrich et al., 2015; Sakuragi et al., 2023; Kim and Budin, 2024; Juarez-Contreras et al., 2025). In this study, we found that genes involved in GPI lipid remodeling and endocytosis are also required for vacuolar Lo domain formation in yeast. In WT cells, remodeled GPI-APs that accumulate in sterol- and sphingolipid-enriched plasma membrane Lo domains may be transported to the vacuole along with these domain-forming lipids. In this case, GPI lipid remodeling may play a role in delivering sterols and sphingolipids to the vacuole to promote vacuolar Lo domain formation. Alternatively, only GPI-APs dissociated from the Lo domain of the plasma membrane due to glucose depletion may be endocytosed and transported to the vacuole. Upon reaching the vacuole, the protein moiety of GPI-APs is degraded, whereas the lipid moiety remains within the vacuolar membrane and functions to form Lo domains. It is unclear whether the remodeled lipid moiety of GPI-APs functions as a carrier that transports sterols and sphingolipids to the vacuole via endocytosis, or plays a role as a component of the vacuolar Lo domain, or both. Future studies are required to determine the precise role of GPI lipid remodeling in vacuolar Lo domain formation.

## Materials and Methods

### Yeast strains and plasmids

All strains of *Saccharomyces cerevisiae* and plasmids used in this study are listed in TableS1. Gene deletion and fluorescent protein tagging were performed using a PCR-based one-step gene replacement method. To construct the Vph1-mCherry expression plasmid, the *VPH1*-mCherry fusion gene was PCR-amplified from genomic DNA of a cell with mCherry tagged to the C-terminus of Vph1 and cloned into the HindIII-XhoI sites of pRS415-TDH3. For the Erg6-GFP expression plasmid, the *ERG6* ORF amplified from genomic DNA and the GFP coding sequence from pGREG600 were cloned into pRS416-TDH3 by In-Fusion Cloning.

### CULTURE conditions and glucose starvation

Yeast strains were cultured at 25 □ in YPD rich medium (1% yeast extract, 2% peptone, 2% glucose, and 0.2% adenine). Synthetic defined (SD) medium (0.17% yeast nitrogen base without amino acids, 0.5% ammonium sulfate, and 2% glucose) supplemented with appropriate amino acids and bases or synthetic complete (SC) medium (0.17% yeast nitrogen base without amino acids, 0.5% ammonium sulfate, 2% glucose, and complete supplement mixture) was used for auxotrophic selection and imaging. For glucose starvation, cells grown to logarithmic phase in the media indicated in the corresponding figure legends were collected by centrifugation (3,000 *g* for 3 min), washed twice and resuspended in SD medium lacking glucose (SD-glucose), and incubated for the times indicated in the figure legends.

### Fluorescence microscopy

Cells were imaged using a fluorescence microscope equipped with differential interference contrast (DIC) optics. LDs were visualized by staining with AutoDot (1 µg/ ml final concentration; Abcepta, San Diego, CA, USA) for 10 min, followed by washing with the corresponding culture medium prior to imaging. The number of LDs per cell, the presence or absence of vacuolar microdomains, and categorical classification of fluorescence patterns were assessed by visual inspection. Fluorescence intensity ratios between the plasma membrane and vacuole for Gas1-GFP were measured using the line profile function in ImageJ (National Institutes of Health, Bethesda, MD, USA). For Gas1-GFP and Cwp2-VENUS visualization analyzed in Fig. S3, cells were cultured under the conditions described in the figure legends, washed once with PBS (pH 7.4), and incubated on ice for 20 min prior to fluorescence imaging.

### TLC analysis of neutral lipids

Yeast cells were grown in SD medium and either analyzed directly or after a 24 h shift to SD-Glucose. Cells were set to an OD_600_ of 10 and treated with a 10 mM NaN_3_-10mM NaF mixture before disruption with glass beads. Total lipids were extracted using chloroform-methanol-water (CMW, 10:10:3, v/v/v) and separated by thin-layer chromatography (TLC) as described previously (Yang et al., in press; ikeda et al., 2020). For the solvent system, petroleum ether–diethyl ether–acetic acid (25:25:1, v/v/v) was used for the first third of the plate, followed by petroleum ether–diethyl ether (49:1, v/v) for the remaining distance. Lipids were visualized by staining with 0.63 g MnCl□·4H□O dissolved in methanol-water-sulfuric acid (60:60:4, v/v/v) for 10 s, then heated at 110 °C for 15 min. Lipid bands were quantified using ImageJ.

### Western blot analysis

Cells expressing Erg6-GFP were cultured under the conditions described in the figure legends and subjected to glucose starvation. Cell lysates denatured with SDS sample buffer were separated by SDS-PAGE (Kajiwara et al., 2008), and analyzed by Western blotting using a mouse monoclonal anti-GFP antibody (clone 7.1 and 13.1, Cat# 11814460001, Roche, Basel, Switzerland) (1:1000, v/v) followed by horseradish peroxidase–conjugated anti-mouse IgG (Cat# A4416, Sigma-Aldrich, St. Louis, MO, USA) (1:20000, v/v). Signals were detected by ECL enhanced chemiluminescence. Erg6-GFP degradation was quantified as the ratio of cleavaged Erg6-GFP band intensity to total Erg6-GFP signal (Erg6-GFP + cleavaged Erg6-GFP).

### Statistical analysis

All statistical analyses were performed using R (R Foundation for Statistical Computing, Vienna, Austria). Comparisons between two groups were performed using a two-tailed Student’s t test, whereas comparisons among multiple groups were performed by one-way ANOVA followed by Tukey’s multiple comparison test, as indicated in the figure legends. Categorical data were analyzed using the Chi-square test. Data are presented as mean ± S.D. unless otherwise stated. The number of independent biological replicates is indicated as *n* in the corresponding figure legends (ns: not significant; *, p<0.05; **, p<0.01; ***, p<0.001). All statistical analyses performed in this study are provided in Table S2.

## Declaration of AI Tool Use in Manuscript Preparation

In this study, ChatGPT (OpenAI, San Francisco, CA, USA) was used to assist with language editing and optimization of R code for graph generation and statistical analyses, including Student’s t test, Chi-square test, and Tukey’s multiple comparison test. All AI-assisted revisions were reviewed and validated by the authors, who take full responsibility for the content of the final manuscript.

## Acknowledgments

We sincerely thank Dr Martin Funk and Dr Laura Popolo for generously providing the plasmids, pRS4XX empty vectors and pRS416-GAS1-GFP, respectively.

## Competing interests

The authors declare no competing or financial interests.

## Author contributions

K.M., K.H., Y.Y., H.N., A. CR., S. LM., and M.M.: Investigation.

K.M. and K.H.: Data curation and formal analysis.

K.M., K.H., M.M. and K.F.: Writing – original draft; writing – review and editing.

K.F.: Funding acquisition, conceptualization, and project administration.

## Funding

This work was supported by the Japan Society for the Promotion of Science (JSPS), Grants-in-Aid for Scientific Research (KAKENHI), Japan (19H02922) to K. F., and Grant PID2023-151267NB-I00 to M.M. funded by MICIU/AEI/ 10.13039/501100011033 and, as appropriate, by “ERDF A way of making Europe”, by “ERDF/EU”, by the “European Union” or by the “European Union NextGenerationEU/PRTR”.

## Notes

### Competing Interest Statement

The authors have declared no competing interest.

