## Supplemental figure for "GPI lipid remodeling regulates lipophagy by forming lipid domains in response to glucose deprivation"

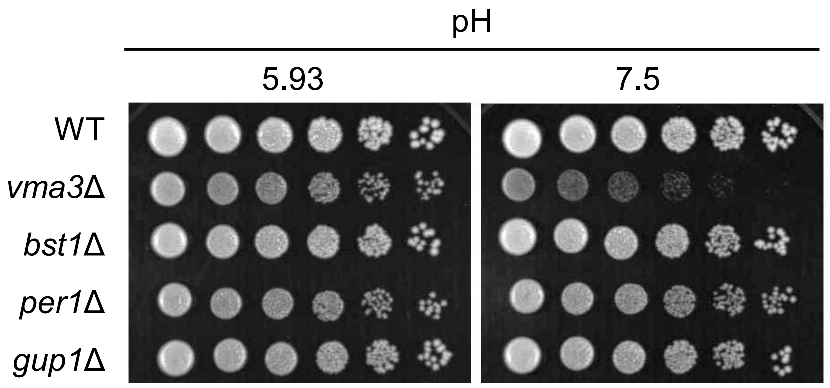


**Fig. S1. GPI remodeling mutants do not exhibit alkaline sensitivity.**
Serial dilution spot assays were performed on YPD plates adjusted to pH 5.9 or pH 7.5. WT, GPI remodeling mutant strains, and the vacuolar ATPase mutant *vma3Δ* were normalized to $OD_{600}$=1, serially diluted (1:5), and spotted onto plates, followed by incubation for 3 days. The *vma3Δ* strain exhibited impaired growth at pH 7.5, whereas GPI remodeling mutants displayed growth comparable to WT.

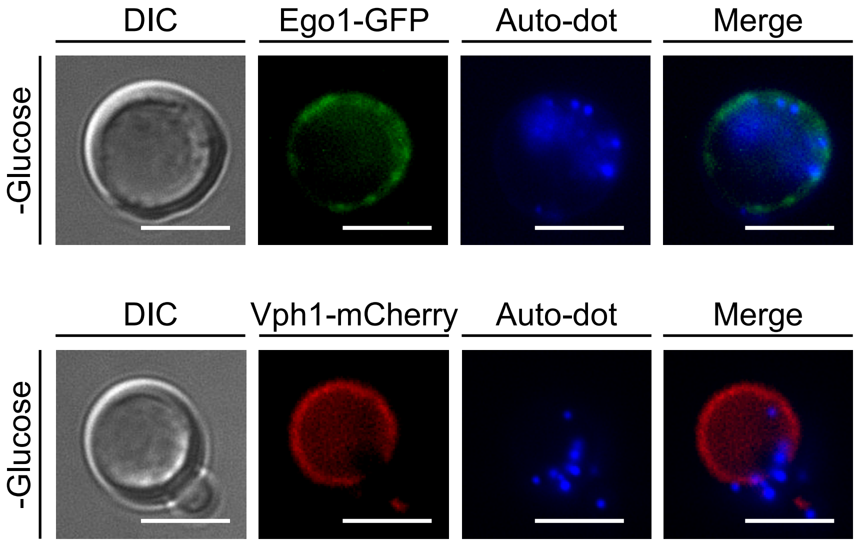


**Fig. S2. LDs accumulate at vacuolar Lo domains under glucose starvation.**
Wild-type (WT) cells expressing Ego1-GFP (Lo domain marker) or Vph1-mCherry (non-Lo domain marker) were grown in SD+Glucose and shifted to SD-Glucose for 24 h. Cells were stained with AutoDot to visualize lipid droplets (LDs) and examined by fluorescence microscopy. Representative images are shown. Scale bar, 5 $\mu$m.

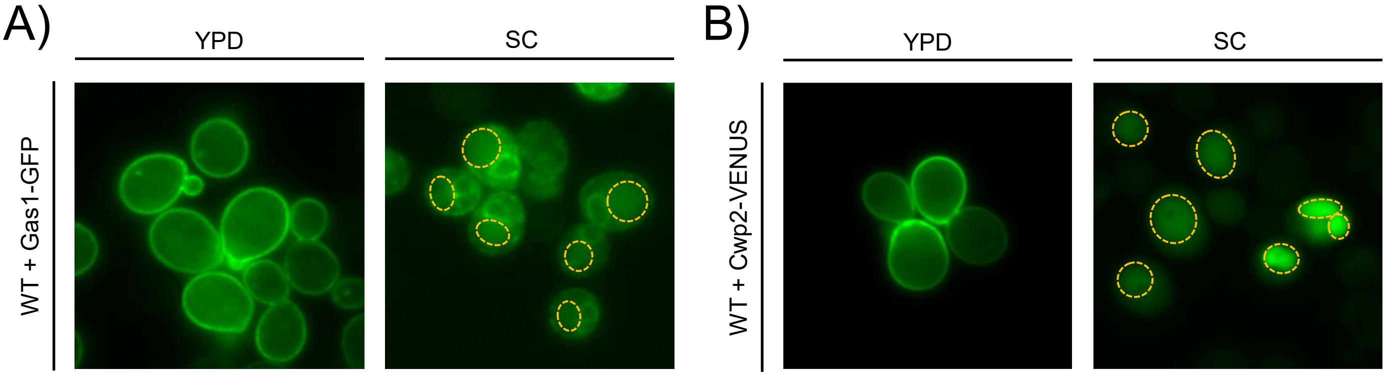


**Fig. S3. GPI-anchored proteins localize to the plasma membrane under nutrient-rich conditions and to the vacuole under nutrient-limited conditions.**
(A), Wild-type (WT) cells expressing Gas1-GFP were grown overnight in either YPD or SC medium containing glucose and visualized by fluorescence microscopy. Representative images are shown. The vacuoler membrane is outlined by yellow dashed lines. Scale bar, 5 μm. (B), Wild-type (WT) cells expressing Cwp2-VENUS were grown overnight in either YPD or SC medium containing glucose and visualized by fluorescence microscopy. Representative images are shown. Scale bar, 5 μm.

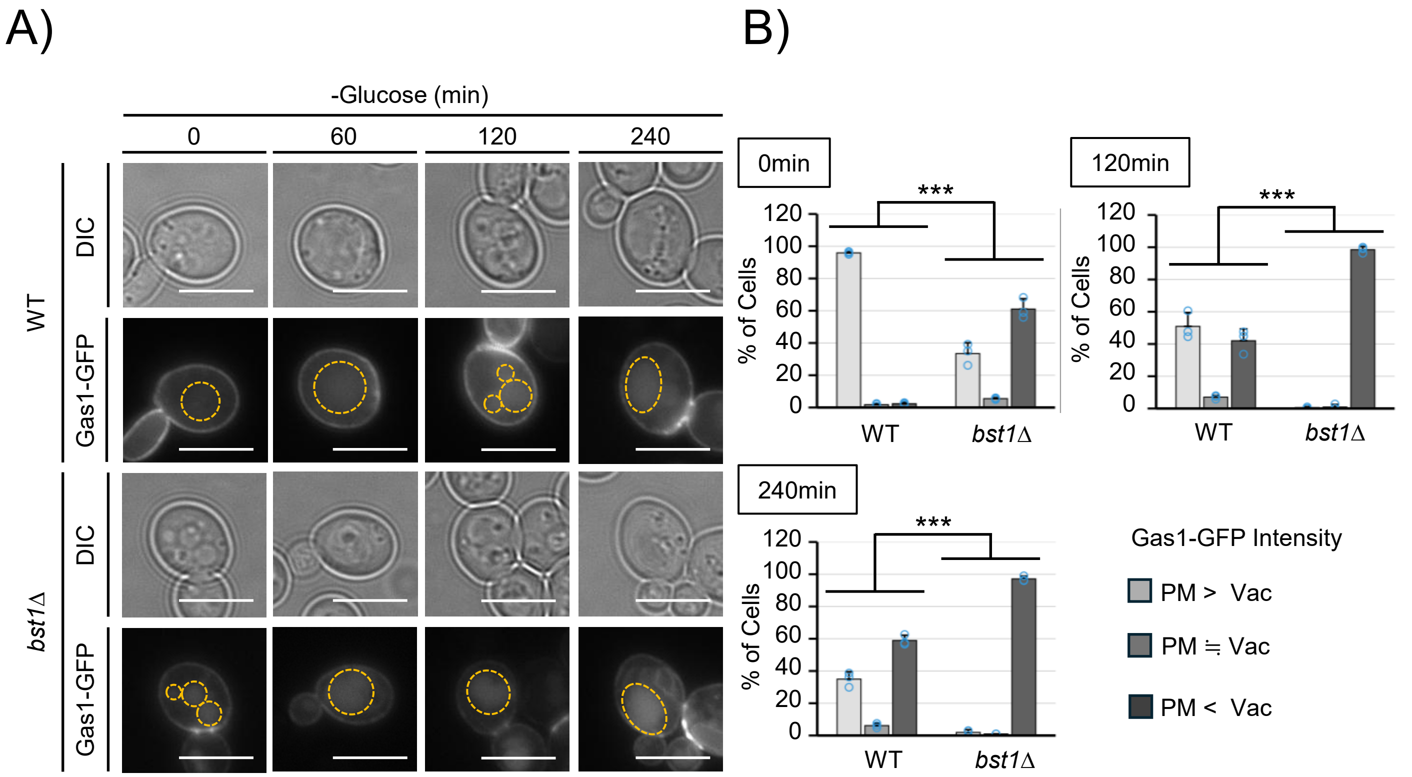


**Fig. S4. Deletion of *BST1* facilitates vacuolar localization of Gas1-GFP during glucose starvation.** (A), WT and *bst1*Δ cells expressing Gas1-GFP were grown and analyzed as described in Fig. 4A. Cells were examined at 0, 60, 120, and 240 min after shifting to SD-Glucose. Representative images are shown. The vacuoler membrane is outlined by yellow dashed lines. Scale bar, 5 μm. (B), Quantification of Gas1-GFP localization was performed as described in Fig. 4B. The percentage of cells in each category (PM > vacuole, PM = vacuole, and PM < vacuole) was determined at 0, 120, and 240 min after shifting to SD-Glucose. ≥100 cells were analyzed per biological replicate. Data represent mean ± S.D. from three independent experiments ($n = 3$). Statistical significance was assessed using a Chi-square test. ***$p < 0.001$.
